# Single-cell spatial transcriptomics of inherited retinal degeneration reveals broad and focal organization of disease responses

**DOI:** 10.64898/2026.09.28.751688

**Authors:** Aya A. Abu-Alfa, Ryan Passino, Zhangyong Wei, Sumasri Kotha, Arjun Raj, Katherine E. Uyhazi

**Affiliations:** Perelman School of Medicine, University of Pennsylvania, Philadelphia, PA 19104, USA; Department of Ophthalmology, Perelman School of Medicine, University of Pennsylvania, Philadelphia, PA 19104, USA; Department of Neuroscience, Perelman School of Medicine, University of Pennsylvania, Philadelphia, PA 19104, USA; Department of Bioengineering, School of Engineering and Applied Science, University of Pennsylvania, Philadelphia, PA 19104, USA; Department of Genetics, Perelman School of Medicine, University of Pennsylvania, Philadelphia, PA 19104, USA

## Abstract

Inherited retinal degenerations are largely monogenic disorders characterized by progressive photoreceptor dysfunction and death. Although the transcriptomic responses to inherited degeneration are well-characterized, how these responses are spatially organized across the retinal axis has not been mapped. Here, we used seqFISH to spatially profile 173 cell identity and degeneration-associated genes with single-cell, single-transcript resolution in male wild-type and *LCA5^gt/gt^*mouse retinas at early, intermediate, and advanced disease stages. In early disease, rod injury and stress response markers were enriched in the superior retina, while rod function and identity markers were enriched in the inferior retina. Nested within the broader superior-inferior organization, a coordinated rod injury response comprising *Edn2*, *Fgf2*, *Socs3*, and *Stat3* was organized into focal regions on the order of 100 μm in width, with high-response regions directly adjacent to low-response regions. These focal regions were locally enriched for markers of Müller reactive gliosis and rod stress responses. Our single-cell spatiotemporal map reveals that transcriptional responses to degeneration vary at both the broad and focal scales, and offers a framework for investigating how local tissue properties and interactions among neighboring cells may contribute to disease progression.

## Introduction

Inherited retinal degenerations are largely monogenic disorders in which photoreceptor dysfunction and death cause progressive vision loss.^1^ Transcriptional responses to degeneration have been well-characterized in photoreceptors, Müller glia, and other retinal cell types.^2–10^ Single-cell studies have begun to show that distinct photoreceptor and glial transcriptional states coexist at the same disease stage, suggesting that cells of the same type can vary in their disease response despite having the same causal mutation.^2,9^ However, whether these responses are spatially organized across the retinal axis has not been mapped. Because tissue dissociation removes spatial information, it is unknown whether cells with different transcriptional states are interspersed across the retina or concentrated in particular regions. Defining the spatial organization of disease responses could point to variation in local tissue conditions and cell-cell interactions as potential contributors to heterogeneity among cells carrying the same mutation.

Spatial transcriptomic approaches have been applied to several mouse models of inherited retinal degeneration, including *rd1*, *Rs1*, *Mfrp*, and *Prph2* mutant retinas.^11–14^ However, these studies used spatial information primarily to characterize transcriptional changes within retinal layers or cell types, rather than to map how these responses vary with retinal position. Evidence that transcriptional responses can vary with retinal position instead comes from models of experimentally induced degeneration. Specifically, following hyperoxic injury, markedly different transcriptional responses were induced in the superior versus inferior retina, while a recent Visium spatial transcriptomics study of photo-oxidative injury identified a highly localized inflammatory and stress response in the superior retina.^15,16^ However, both models rely on degeneration induced by supraphysiologic insults, which is biologically distinct from progressive degeneration caused by an inherited mutation. Therefore, while these studies suggest that degeneration-associated transcriptional responses can vary with broad retinal position, whether and at what scale such responses are spatially organized in inherited retinal degeneration remains unknown. Addressing this gap requires simultaneous measurement of cell identity, gene expression, and position in the intact degenerating retina.

Here, we applied sequential fluorescence *in situ* hybridization (seqFISH) to map degeneration-associated transcriptional responses in the *LCA5^gt/gt^* mouse retina at early, intermediate, and advanced disease stages. In early disease, injury and stress-associated transcription was greater in the superior retina, whereas rod function and identity transcripts were relatively enriched inferiorly. Within the broad pattern, injury-associated transcription in rods was organized into repeated focal regions enriched for markers of Müller reactive gliosis and rod stress responses. Our spatial transcriptomic map shows that, within a shared genetic background, degeneration-associated transcriptional responses in the retina spatially vary at both the broad and focal scales.

## Results

### seqFISH resolves major cell types and the superior-inferior retinal axis

To map transcriptional responses across the degenerating retina, we used a model of degeneration with reproducible photoreceptor loss. The *LCA5^gt/gt^*mouse model contains a homozygous gene-trap insertion that results in loss of functional lebercilin (LCA5), a ciliary protein required for trafficking of photoreceptor proteins from the inner to the outer segment.^17,18^ Loss of lebercilin disrupts outer segment formation and photoreceptor protein localization, followed by photoreceptor death, although the mechanisms leading to photoreceptor death remain incompletely understood.^17,18^ The *LCA5^gt/gt^*mouse undergoes rapid, progressive photoreceptor loss that recapitulates that seen in patients with homozygous *LCA5* mutations.^17,19^ Because defects in photoreceptor ciliary structure and trafficking underlie many inherited retinal degenerations, the *LCA5^gt/gt^*model is relevant to the broader class of retinal ciliopathies.^20,21^ Its predictable disease course^17,19^ allowed us to sample distinct stages of degeneration within one genetic background.

We focused our spatial analysis on genes whose expression is recurrently altered across genetically distinct models of inherited retinal degeneration. We designed a targeted gene panel of 173 genes informed by published transcriptomic studies across multiple models of inherited retinal degeneration, including *rd1*,^3^ *rd10*,^2,4^ *Prom1*-deficient,^5^ and *VPP* models.^22^ The panel included retinal cell type markers^23–25^ and genes representing recurrent changes in photoreceptor identity and function, ciliary and synaptic organization, injury and stress responses, regulated cell death, Müller glial reactivity, and immune activation (Supplementary Table 1). The resulting panel was therefore designed to capture responses recurring across retinal degeneration models rather than changes specific to *LCA5^gt/gt^*disease.

Using seqFISH with the 173-gene panel, we profiled *LCA5^gt/gt^*retinas at 21, 30, and 64 days of age, corresponding to early, intermediate, and advanced degeneration (Fig. 1a).^19^ We selected 21 and 30 days to capture active transcriptional responses to degeneration while substantial photoreceptor populations remained, and included retinas at 64 days to capture the advanced stage of photoreceptor loss and retinal remodeling. We also profiled wild-type retinas at 21 and 64 days to distinguish degeneration-associated changes from age-related differences. We profiled two biological replicates per condition, each consisting of one section from a separate animal (*n* = 2 animals per condition). Across 10 retinal sections, we detected an average of approximately 5.8 million RNA transcripts per retina, with an average of 324 transcripts and 56 unique genes per segmented cell (Supplementary Fig. 1a).

**Fig. 1:**
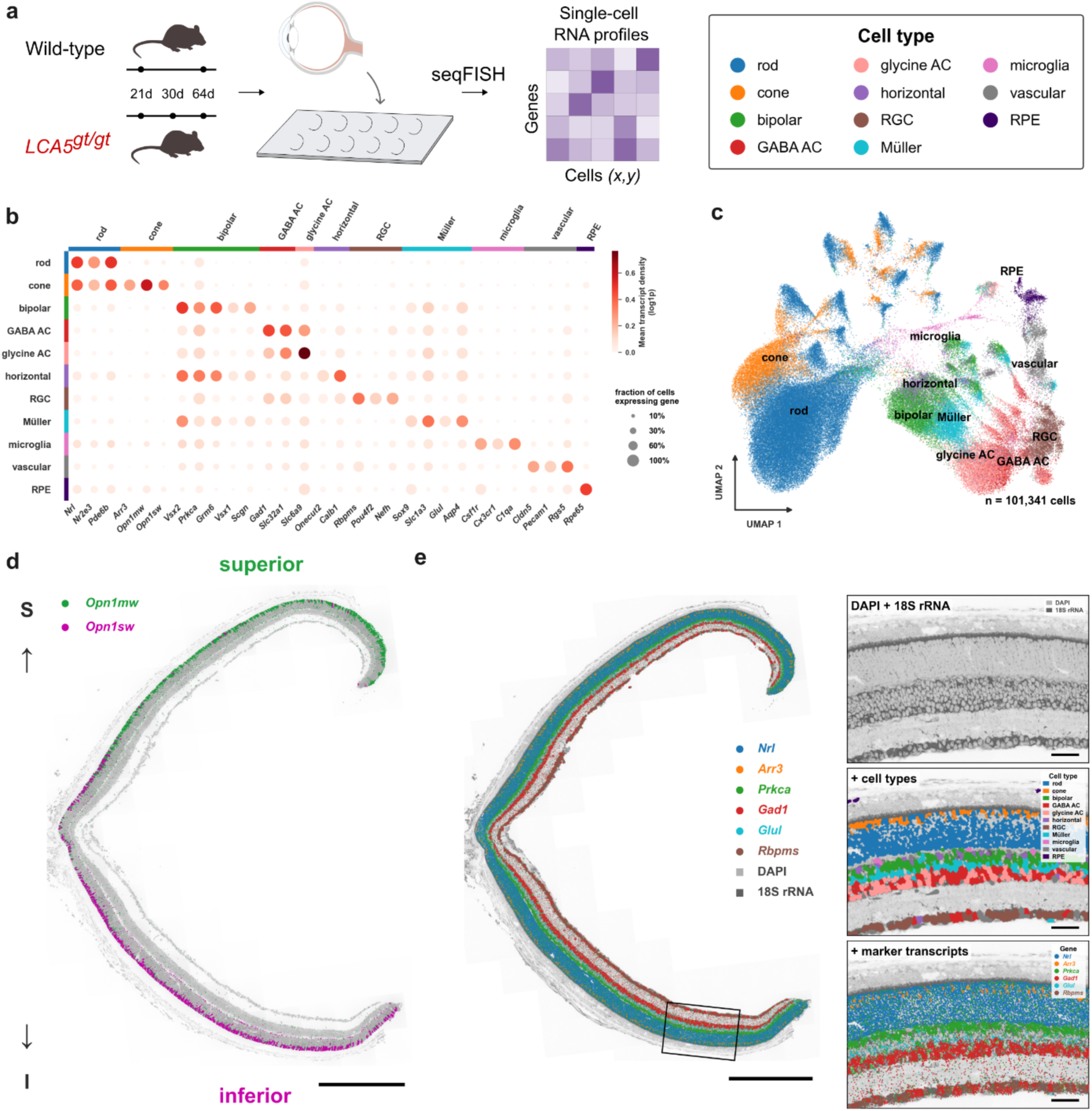
seqFISH resolves major cell types and the superior-inferior retinal axis. **(a)** Schematic of the experimental design and seqFISH workflow. Retinal sections from wild-type mice at 21 and 64 days and *LCA5^gt/gt^* mice at 21, 30, and 64 days were profiled using a targeted panel of 173 genes. Eye illustration was modified from Servier Medical Art, licensed under CC BY 4.0. **(b)** Expression of canonical marker genes across the 11 identified cell types. Dot size indicates the fraction of cells expressing each gene, and color indicates mean log_1_p-transformed transcript density per cell within each cell type. **(c)** Harmony-integrated Uniform Manifold Approximation and Projection (UMAP) embedding of 101,341 typed cell bodies pooled across conditions and biological replicates, colored by cell type. 11 major cell types were identified: rods, cones, bipolar cells, GABAergic amacrine cells (GABA AC), glycinergic amacrine cells (glycine AC), horizontal cells, retinal ganglion cells (RGC), Müller glia, microglia, vascular cells, and retinal pigment epithelium (RPE). **(d)** Representative wild-type retinal section at 21 days showing *Opn1mw* and *Opn1sw* transcripts used to orient the superior-inferior retinal axis. Scale bar, 500 μm. **(e)** Left, representative wild-type retinal section at 21 days showing several cell type marker transcripts. Box indicates the magnified region. Right, magnified region showing DAPI and 18S rRNA staining, segmented cells colored by assigned cell type, and cell type marker transcripts. Scale bars, 500 μm for the full retinal section and 50 μm for the magnified region.

Using expression of canonical marker genes, we assigned 101,341 cell bodies to 11 major cell types: rods, cones, bipolar cells, GABAergic and glycinergic amacrine cells, horizontal cells, retinal ganglion cells, Müller glia, microglia, vascular cells, and retinal pigment epithelium (Fig. 1b, c). Cell bodies assigned to each cell type occupied the expected retinal layers, providing an independent validation of the cell type assignments (Supplementary Fig. 1b). The relative abundance of cell types and the mean cellular transcript density of each gene showed strong correlations between biological replicates in each condition, with a median Pearson’s *r* of approximately 0.99 for both metrics (Supplementary Fig. 1c). Because transcripts in Müller glia fibers, neuronal processes, and photoreceptor inner and outer segments could not be confidently assigned to individual cell bodies, these transcripts were excluded from cell body-level analyses but retained for tissue- and layer-level analyses.

Because prior retinal injury studies have shown differences in transcriptional responses in the superior versus inferior retina,^15,16^ we chose to orient retinal sections along the superior-inferior axis. To do so, in addition to preserving orientation during tissue preparation (see Methods), we used the well-established reciprocal expression gradients of the cone opsins *Opn1mw* and *Opn1sw*^26^ (Fig. 1d). As expected, *Opn1mw* was enriched toward the superior retina, whereas *Opn1sw* was enriched toward the inferior retina. The opposing gradients remained detectable across all genotypes and ages (Supplementary Fig. 1d), allowing sections to be oriented consistently despite progressive degeneration. The expected superior enrichment of *Aldh1a1*^27^ independently supported the assigned orientation in all conditions (Supplementary Fig. 1d). A representative wild-type retinal section at 21 days illustrates the multiple layers of information captured by seqFISH, including tissue morphology defined by DAPI and 18S rRNA staining, cell type assignments, and the spatial positions of individual transcripts for selected marker genes (Fig. 1e).

Together, these data provide spatially resolved maps of major cell types and expression of 173 genes across the superior-inferior axis in wild-type and degenerating *LCA5^gt/gt^*retinas. We next used these maps to compare how cell type composition and gene expression varied across the retina with progressive degeneration.

### seqFISH resolves progressive retinal remodeling across *LCA5^gt/gt^*degeneration

To characterize retinal remodeling in degeneration, we quantified changes in retinal structure and cellular composition over the course of *LCA5^gt/gt^* degeneration. Compared to wild-type, there was an early reduction in both the absolute number and relative abundance of rods in *LCA5^gt/gt^* retinas at 21 days, followed by further decline at 30 days and near depletion by 64 days (Fig. 2a-d; Supplementary Fig. 2b). In contrast, the absolute number of cones was relatively similar at 21 days and 30 days before declining at 64 days (Supplementary Fig. 2b). Because rods were preferentially lost, the relative abundance of cones increased with progressive degeneration, and cones comprised most of the surviving photoreceptors at 64 days (Fig. 2a-d; Supplementary Fig. 2a). Furthermore, the relative abundance of microglia was increased across *LCA5^gt/gt^* retinas compared to wild-type retinas (Fig. 2a-d) and microglia were observed in the outer nuclear layer in *LCA5^gt/gt^*retinas at 21 and 30 days (Fig. 2c), consistent with previous studies showing microglial proliferation and recruitment in inherited retinal degenerations.^28–30^ The outer nuclear layer progressively thinned over the course of degeneration, accompanied by disruption of photoreceptor inner and outer segment structure (Fig. 2b, c, e). In contrast, the inner nuclear layer thickness was preserved, and inner retinal cell types remained represented throughout degeneration (Fig. 2a-e). The relative abundance of rods was also spatially asymmetric, with a lower relative abundance in the superior than inferior *LCA5^gt/gt^* retina at 21 and 30 days (mean superior to inferior ratios of 0.84 and 0.87, respectively, Supplementary Fig. 2c).

**Fig. 2:**
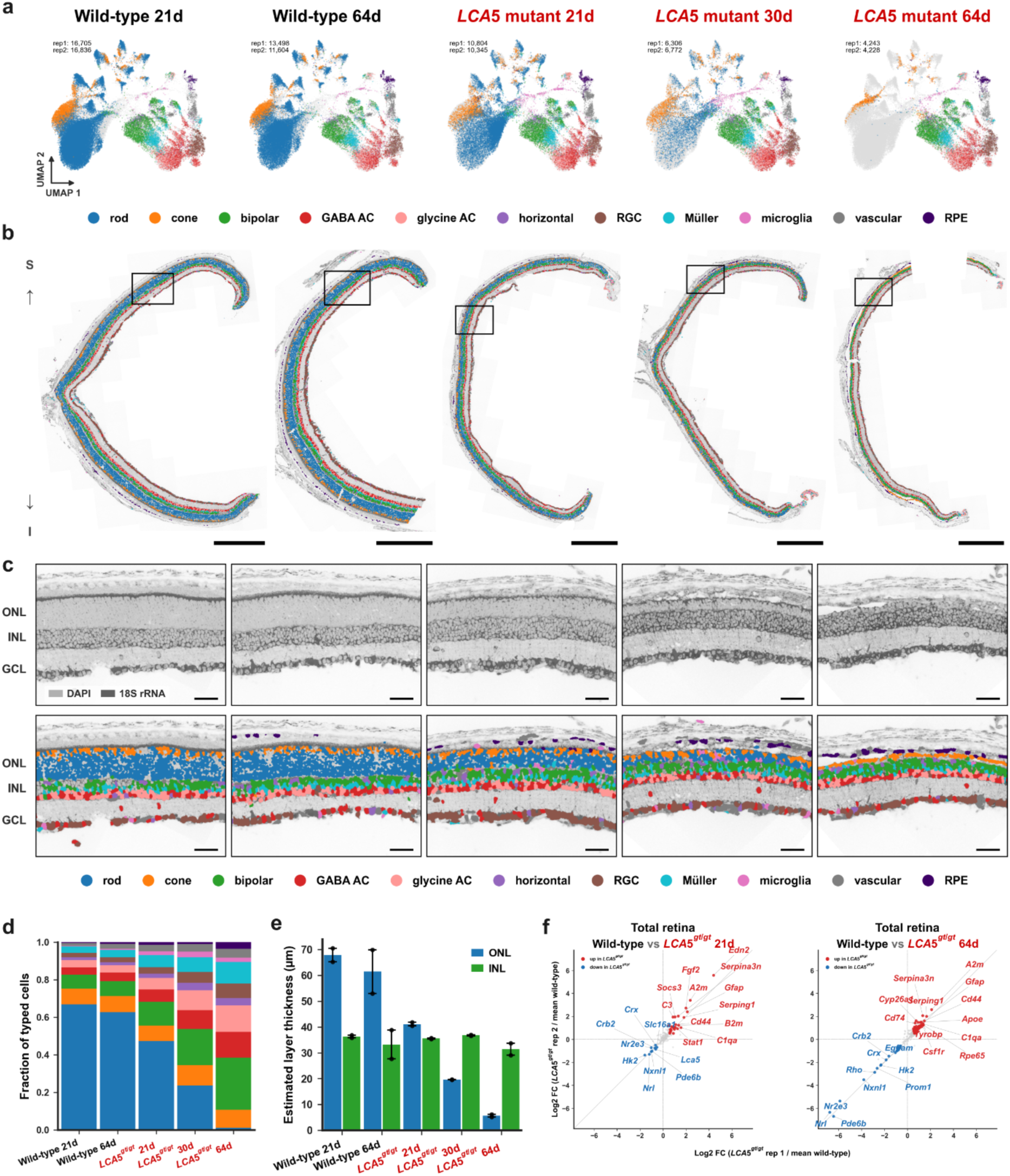
seqFISH resolves progressive retinal remodeling across *LCA5^gt/gt^* degeneration. **(a)** UMAP embedding of all typed cells. Cells pooled from each condition are colored by assigned type. The number of typed cells in each replicate is shown. **(b)** Representative full retinal sections from each condition, oriented along the superior-inferior axis, with cells colored by assigned type. Boxes indicate the regions magnified in (c). Scale bars, 500 μm. **(c)** Magnified regions indicated in (b). Top, DAPI and 18S rRNA staining with the outer nuclear layer (ONL), inner nuclear layer (INL), and ganglion cell layer (GCL) labeled. Bottom, cells colored by assigned type. Scale bars, 50 μm. **(d)** Cell type composition per condition, expressed as the fraction of typed cells assigned to each type. Stacked bars show the mean of individual replicates. **(e)** Estimated outer nuclear layer (ONL) and inner nuclear layer (INL) thicknesses across conditions. Points represent individual replicates; bars show the mean; whiskers span the minimum and maximum replicate values. **(f)** Replicate-concordant transcriptional changes in relative transcript abundance across the total retina between wild-type and *LCA5^gt/gt^*retinas at 21 (left) and 64 (right) days. Axes show log₂-fold changes for each replicate. Increased genes are shown in red, decreased genes in blue, and all other genes in gray.

To measure tissue-level transcriptional changes associated with degeneration, we compared relative transcript abundance between *LCA5^gt/gt^* and age-matched wild-type retinas at 21 and 64 days. At 21 days, before complete collapse of the outer nuclear layer, *LCA5^gt/gt^*retinas already showed consistent increases in injury response transcripts (*Edn2, Fgf2, Socs3*)^2,31,32^ and Müller reactive gliosis *(Gfap*, *Serpina3n, Cd44*)^8,9,33,34^ (Fig. 2f). By 64 days, when the outer nuclear layer had collapsed, the relative abundance of rod identity and function transcripts such as *Nrl*, *Nr2e3*, *Pde6b*, and *Nxnl1*^24^ was markedly reduced, whereas several Müller reactive gliosis and inflammatory transcripts remained elevated (Fig. 2f).

Our findings recapitulate the progressive collapse of the outer nuclear layer, loss of photoreceptor outer segments, and preservation of the inner retina observed over the course of *LCA5^gt/gt^* degeneration.^17,19^ Notably, we identify substantial preservation of cone photoreceptors in advanced-stage *LCA5^gt/gt^* mice, a feature that has not, to our knowledge, been previously described at this stage of degeneration. We further show that the *LCA5^gt/gt^* retina engages several injury, glial, and inflammatory transcriptional responses reported across inherited retinal degeneration models. Because the tissue-level measurements are influenced by changes in cellular composition, we next focused on transcriptional responses within surviving rods and their spatial distribution across the degenerating retina.

### Early transcriptional responses to degeneration in rods show superior-inferior asymmetry

Having established the superior-inferior orientation of each retinal section, we next asked whether degeneration-associated transcriptional responses within surviving rods varied along this axis. We focused the analysis on *LCA5^gt/gt^* retinas at 21 days, when sufficient rods remained for comparison and age-matched wild-type controls were available.

To identify genes with reliable rod expression, we applied a filter requiring expression above the estimated spillover signal from neighboring cells and extended cellular processes, such as Müller glia fibers (see Methods). We retained 56 genes for analysis (Supplementary Fig. 3a), of which 22 met effect size and replicate concordance criteria in *LCA5^gt/gt^*rods at 21 days relative to wild-type rods at 21 days (Fig. 3a). 17 genes were increased, including *Edn2*, a photoreceptor injury signal induced across inherited and experimentally induced retinal damage models, *Fgf2*, *Socs3*, and *Stat3*, which are associated with the leukemia inhibitory factor/JAK-STAT3 injury response, and the photoreceptor stress response genes *Cebpd*, *Gadd45b*, and *Nr4a1*.^2,6,31,32,35^ *Rho* and *Prph2* transcripts were also increased, which may be related to disrupted outer segment formation, although mislocalization of RHO in *LCA5^gt/gt^* photoreceptor nuclei has been shown at the protein rather than the transcript level.^17^ 5 genes were reduced, including the rod identity factor *Nrl*, the metabolic genes *Hk2* and *Slc16a1*, and the trophic support gene *Nxnl1*.^24,36–38^ Early degeneration was therefore associated with strong induction of injury and stress-associated transcription and reduced expression of several genes associated with rod identity and function.

**Fig. 3:**
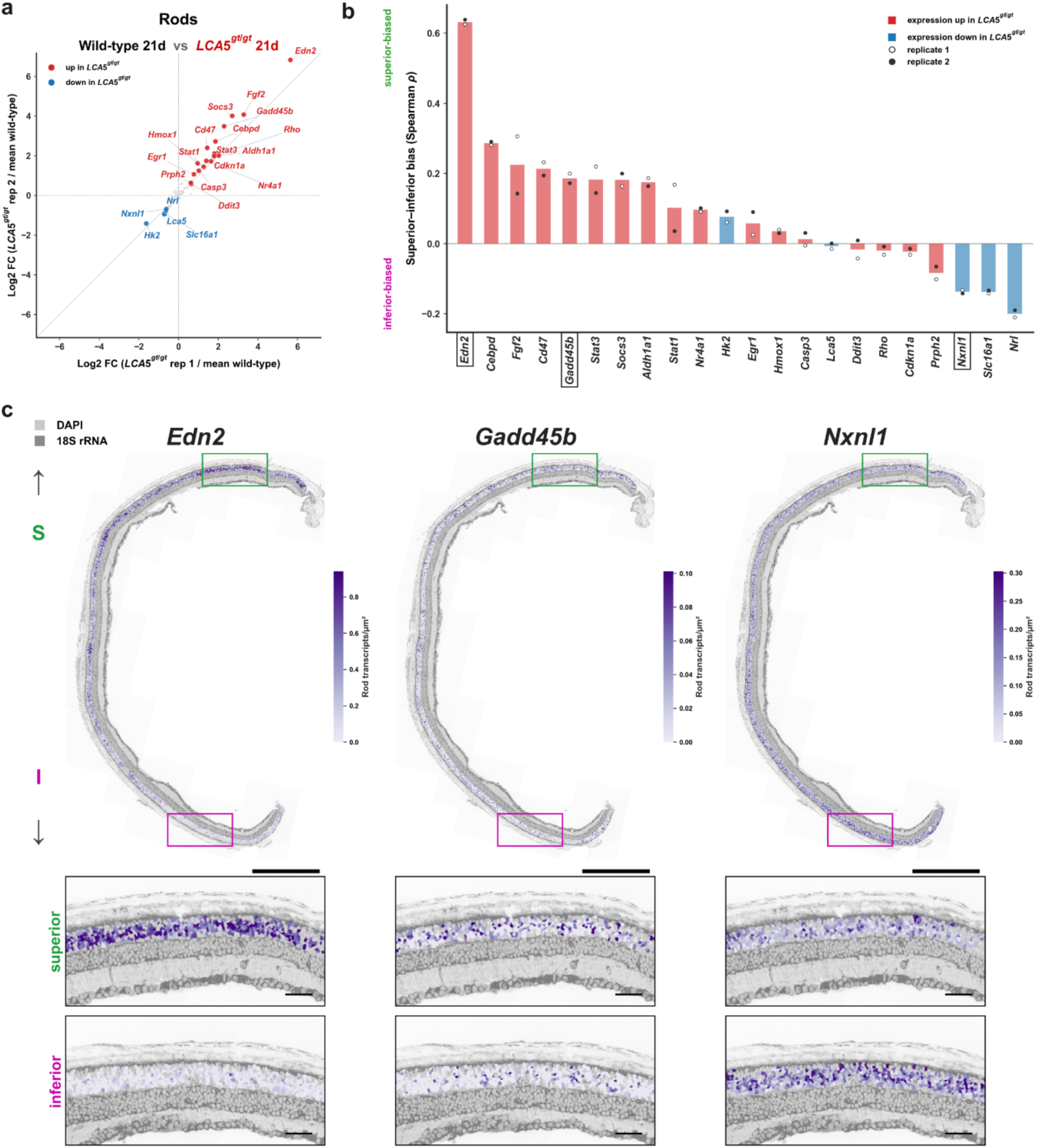
Early transcriptional responses to degeneration in rods show superior-inferior asymmetry. **(a)** Replicate-concordant transcriptional changes in *LCA5^gt/gt^* rods at 21 days compared to age-matched wild-type rods. Axes show log₂-fold changes for each replicate. Increased genes are shown in red, decreased genes in blue, and all other genes in gray. **(b)** Superior-inferior spatial distribution of the 22 genes identified in (a). Bars show the mean Spearman correlation (ρ) between rod transcript density and the smoothed cone opsin superior-inferior gradient across two replicates; open and filled circles represent individual replicates. **(c)** *Edn2*, *Gadd45b*, and *Nxnl1* expression in rods in a representative *LCA5^gt/gt^* retinal section at 21 days. Boxed regions from the superior and inferior hemispheres are magnified below. Each gene is shown using its own scale. Scale bars, 500 μm for full sections and 50 μm for magnified regions.

We then quantified whether these early transcriptional changes in rods varied along the superior-inferior retinal axis. Within each retina, we used the Spearman rank correlation to quantify the relationship between rod expression of each gene and rod position along the superior-inferior axis. Because many injury response transcripts were lowly expressed in wild-type rods, we performed the primary spatial analysis in *LCA5^gt/gt^* rods at 21 days. For context, the same genes showed little replicate-concordant variation along the superior-inferior axis in age-matched wild-type rods (Supplementary Fig. 3b).

*Edn2* showed the strongest replicate-concordant enrichment toward the superior retina. *Cebpd, Socs3, Fgf2, Stat3, Gadd45b,* and *Nr4a1* showed more modest but reproducible superior biases (Fig. 3b, c). In contrast, *Nrl, Slc16a1,* and *Nxnl1* were relatively enriched toward the inferior retina (Fig. 3b, c). Therefore, several injury- and stress-associated transcripts were enriched superiorly, whereas transcripts associated with rod identity and function were relatively enriched inferiorly in early degeneration.

### A rod injury response exhibits focal spatial organization

In addition to the broad superior-inferior differences described above, we observed substantial variation in rod transcript expression at a finer spatial scale. We first focused on *Edn2, Fgf2, Socs3,* and *Stat3*, which were among the most highly induced transcripts in *LCA5^gt/gt^*rods (Fig. 3a, Supplementary Fig. 4a) and are known to be co-induced across multiple models of retinal degeneration, including light damage, VPP, *cpfl1*, and *rd10* as part of the recurrent JAK/STAT3 retinal injury response.^2,32,35,39^ We summarized their expression into an injury response score for each rod (Fig. 4a). In *LCA5^gt/gt^* retinas at 21 and 30 days, the rod injury response score had a broad distribution (Fig. 4a), indicating substantial heterogeneity among surviving rods. By 64 days, the few surviving rods did not show comparable injury response scores to those at 21 and 30 days (Fig. 4a). We excluded *LCA5^gt/gt^* retinas at 64 days from subsequent spatial analyses because too few rods remained for reliable quantitative analysis.

**Fig. 4:**
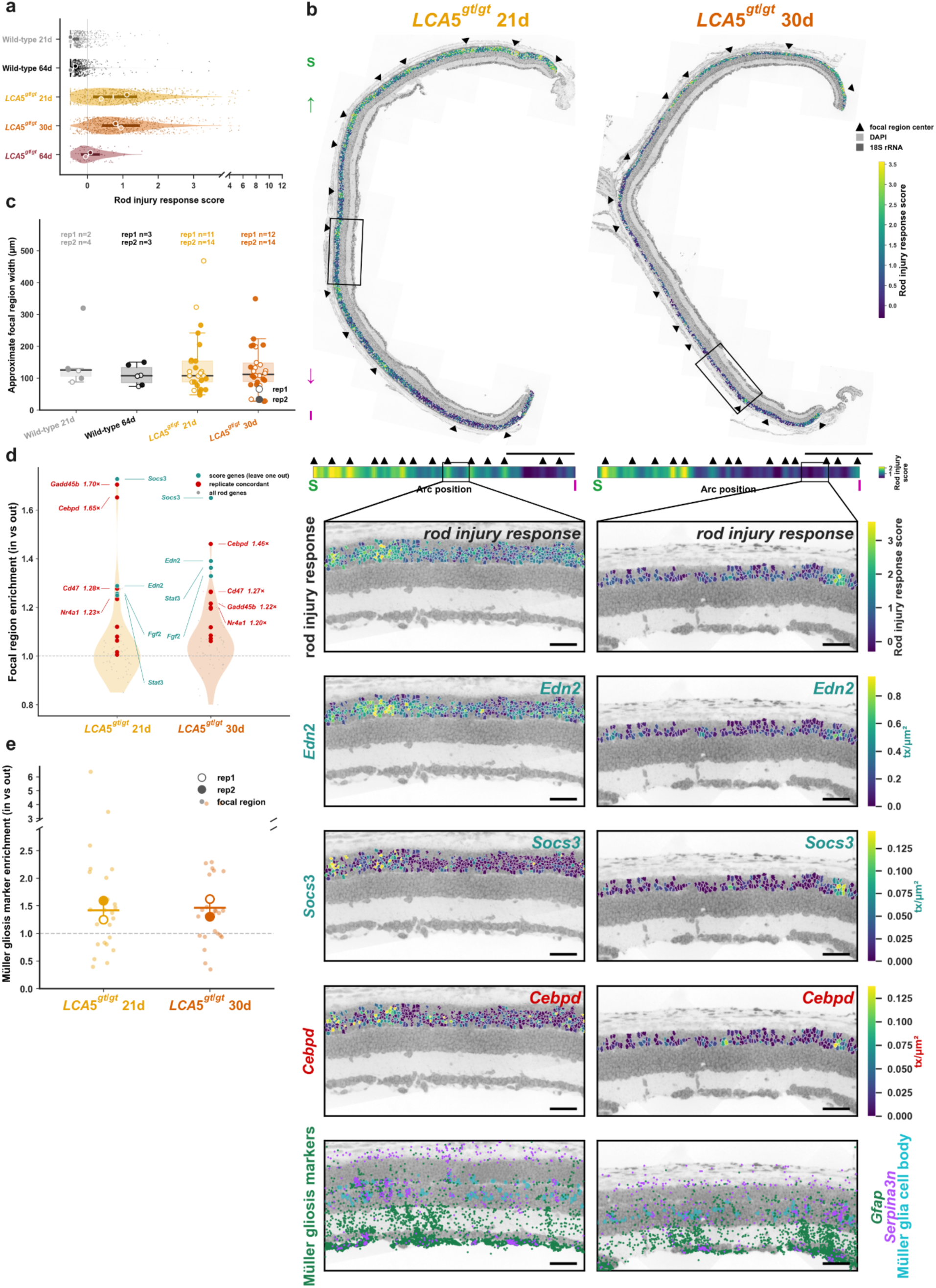
A rod injury response exhibits focal spatial organization. **(a)** Distribution of the injury response score per rod in wild-type retinas at 21 and 64 days and *LCA5^gt/gt^* retinas at 21, 30, and 64 days. Points show a random subsample of up to 500 rods per replicate; circles represent the median of individual replicates; shaded bars show the pooled interquartile range; and white lines indicate the pooled median. **(b)** Top, rod injury response scores mapped onto representative *LCA5^gt/gt^* retinal sections at 21 and 30 days. Arrowheads indicate focal region centers; boxes indicate regions magnified below. Scale bars, 500 μm. Horizontal strips show mean rod injury response score in 25 μm columns along the retinal arc. Bottom, magnified regions showing rod injury response score; rod expression of *Edn2, Socs3,* and *Cebpd*; and Müller reactive gliosis markers *Gfap* and *Serpina3n* with Müller glia cell bodies shown. Scale bars, 50 μm. **(c)** Estimated widths of rod injury response focal regions in wild-type retinas at 21 and 64 days and *LCA5^gt/gt^*retinas at 21 and 30 days. Points represent individual focal regions. The number of focal regions in each replicate is labeled. **(d)** Enrichment of rod transcripts in focal regions relative to the immediately flanking retina in *LCA5^gt/gt^* retinas at 21 and 30 days. Points represent the mean enrichment across individual replicates at each timepoint. Gray points represent all tested rod genes; red points represent genes with an enrichment greater than 1 in each of the four retinas; labeled red points represent genes with a mean enrichment of at least 1.2 across the four retinas; teal points represent the four injury response score genes when tested using focal regions defined by only the other three score genes. **(e)** Enrichment of the Müller gliosis markers *Gfap* and *Serpina3n* in the inner and outer nuclear layers in focal rod injury response regions relative to the immediately flanking retina in *LCA5^gt/gt^* retinas at 21 and 30 days. Points represent individual focal regions; open and filled circles represent pooled region values for individual replicates.

We next mapped the rod injury response score across the retinal sections to assess its spatial organization. Consistent with the analyses of each gene in Fig. 3, the rod injury response score was higher overall in the superior than inferior retina at both 21 and 30 days in *LCA5^gt/gt^* retinas (Supplementary Fig. 4b). Superimposed on this broad difference, however, we observed focal regions of high-scoring rods directly adjacent to comparatively low-scoring rods in both the superior and inferior retina (Fig. 4b). The focal variation was also evident in the spatial expression patterns of the individual injury response score genes (Fig. 4b; Supplementary Fig. 5b).

To quantify the number and spatial scale of the focal regions, we identified local peaks in the rod injury response score profile across the retina and estimated the width of the corresponding regions (see Methods). The two *LCA5^gt/gt^* retinas contained 11 and 14 focal regions at 21 days and 12 and 14 regions at 30 days, compared with 2 and 4 regions in wild-type retinas at 21 days and 3 and 3 regions at 64 days (Fig. 4c). The width of the focal regions typically ranged from approximately 50 to 300 μm, with median widths of approximately 107 μm at 21 days and 112 μm at 30 days (Fig. 4c), indicating a similar spatial scale at both stages. The number of detected focal regions was relatively stable and remained consistently higher in *LCA5^gt/gt^* than in wild-type retinas across a range of peak calling thresholds (Supplementary Fig. 4c). Furthermore, the focal spatial pattern was not dependent on any single score gene: omitting each gene in turn produced a spatial profile that was highly correlated with that produced by the full score (*r* ≥ 0.88; Supplementary Fig. 4d). When focal regions were defined by only three of the four score genes, the omitted fourth gene remained enriched in those regions (Fig. 4d), indicating that all four genes contributed to the focal pattern.

The few focal regions detected in wild-type retinas occurred over a similar spatial scale to those in *LCA5^gt/gt^* retinas, but had substantially lower injury response scores (Fig. 4a, c; Supplementary Fig. 5a). Therefore, focal variation in the rod injury response program was not unique to the degenerating retina, but degenerating retinas were characterized by a larger number of focal regions and a stronger injury response magnitude. These data suggest that degeneration amplifies a tendency toward focal spatial organization of the rod injury response rather than generating focality *de novo*, although they do not establish that the same mechanism generates focal variation in wild-type and *LCA5^gt/gt^* retinas. Across both wild-type and *LCA5^gt/gt^* retinas, the mean rod abundance did not consistently differ within focal regions relative to immediately flanking tissue (Supplementary Fig. 4e).

We next asked whether expression of other degeneration-associated transcripts in rods was locally enriched in these focal regions. Several stress-associated transcripts, namely, *Cebpd, Gadd45b,* and *Nr4a1*, were reproducibly enriched in focal regions relative to immediately flanking tissue in *LCA5^gt/gt^* retinas at 21 and 30 days (Fig. 4b, d; Supplementary Fig. 5b). *Cebpd* is a recurrent photoreceptor injury-associated transcription factor, whereas *Gadd45b* and *Nr4a1* are associated with stress and immediate-early responses, respectively, during retinal degeneration.^2^ *Cd47*, which encodes an anti-phagocytic surface protein,^40^ was also reproducibly enriched in the focal regions. In contrast, *Egr1*, although induced in *LCA5^gt/gt^*rods at 21 days (Fig. 3a), was not enriched in the focal regions. Most other tested rod genes were not consistently enriched in focal regions (Fig. 4d), arguing against the notion that focal regions simply reflect a general increase in transcription or RNA detection. Therefore, the focal rod injury response regions coincided with expression of specific well-established markers of rod stress.

Because Müller reactive gliosis is a major response to photoreceptor degeneration,^8,9,34^ we next asked whether it coincided with the focal rod injury response regions. We focused on *Gfap* and *Serpina3n*, established markers of Müller reactive gliosis that have been experimentally shown to be induced in rat Müller glia following EDN2 exposure.^31,33^ Because Müller cell bodies reside in the inner nuclear layer while their radial processes extend through the outer nuclear layer, we quantified *Gfap* and *Serpina3n* across both layers. The combined *Gfap* and *Serpina3n* transcript density was strongly elevated in *LCA5^gt/gt^* retinas at 21 and 30 days compared to wild-type (Supplementary Fig. 4f) and was higher overall in focal rod injury response regions than in immediately flanking tissue at both stages (Fig. 4b, e). This local enrichment was not solely explained by differences in Müller cell body density, which correlated only weakly with local *Gfap* and *Serpina3n* transcript density (Supplementary Fig. 4g). In contrast, although microglia are recruited to the outer nuclear layer in several inherited retinal degeneration models^28–30^ and were similarly enriched in the outer nuclear layer of *LCA5^gt/gt^* retinas at all disease stages (Supplementary Fig. 4h), their abundance was not consistently altered within the focal regions (Supplementary Fig. 4i).

Together, these analyses show that a rod injury response comprising *Edn2*, *Fgf2*, *Socs3*, and *Stat3* is organized into focal regions during early and intermediate *LCA5^gt/gt^* retinal degeneration. These regions were typically on the order of 100 μm in width, occurred within both the superior and inferior retina, and showed selective enrichment of rod stress responses together with local Müller reactive gliosis. Therefore, the broad superior-inferior gradient captures only one level of spatial variation in the degenerating *LCA5^gt/gt^* retina: within each hemisphere, neighboring populations of surviving mutant rods occupy markedly different injury and stress response states, accompanied by corresponding differences in the transcriptional state of local Müller glia.

## Discussion

In this work, we generated a targeted single-cell spatial transcriptomic map of *LCA5^gt/gt^* retinal degeneration and found that established transcriptional responses are organized at both the broad and focal spatial scales. Rod injury and stress response markers were generally enriched toward the superior retina, whereas rod identity and function markers were relatively enriched inferiorly. Superimposed on this broad organization, rod injury-associated transcription was organized into focal regions on the order of 100 μm in width that were enriched for rod stress response transcripts and markers of Müller reactive gliosis. Rods carrying the same causal mutation therefore differed transcriptionally according to both broad and local anatomical position. The principal advance is therefore not the identification of new disease responses, but the demonstration that established responses have a previously unresolved spatial organization.

At the broad scale, the injury and stress response was strongest in the superior *LCA5^gt/gt^* retina at 21 and 30 days, where rod relative abundance was approximately 15% lower on average than in the inferior retina (Supplementary Fig. 2c). Broad asymmetry in photoreceptor loss is observed in both human inherited retinal degenerations and animal models, but its direction varies with the causal mutation and experimental context.^1,41–44^ Our findings suggest that this broad asymmetry extends to the state of surviving rods. The concordance between lower rod relative abundance and a stronger injury response in the superior retina may reflect broad differences in disease susceptibility, with rods in the superior retina potentially further along the degenerative trajectory than rods in the inferior retina. However, because each timepoint captures only surviving cells, we cannot distinguish this possibility from preferential survival of a more resistant rod subpopulation in the superior retina.

At the focal scale, rods with strong injury-associated transcription were directly adjacent to rods with weaker responses, indicating that transcriptional state can diverge independently of broad anatomical gradients. The enrichment of *Cebpd, Gadd45b,* and *Nr4a1* within these regions further suggests that they represent a broader degeneration-associated stress state. These genes recur in photoreceptor responses across inherited and induced degeneration, although their direct effects on photoreceptor fate are less well established.^2,6,35^ The co-localization of rod injury-associated transcription with markers of Müller reactive gliosis is also consistent with the established photoreceptor-Müller endothelin response, in which EDN2 exposure directly induces *Gfap* and *Serpina3n* expression in cultured Müller glia.^5,31–33^ The focal rod injury response regions may therefore reflect a coordinated multicellular state rather than isolated variation among individual rods. Consistent with a role for coordinated local signaling in producing the focal regions, a previous study showed that leukemia inhibitory factor, which is required for induction of *Edn2, Fgf2, Socs3, Stat3,* and *Gfap* in the VPP retina, is induced in only a subset of Müller glia and proposed that these cells participate in a positive feedback loop with photoreceptors.^32^ Overall, the focal spatial organization suggests that responses to degeneration are shaped not only by genotype but also by shared local factors, such as intercellular signaling, that could contribute to heterogeneity in the vulnerability of genetically identical photoreceptors.

In contrast to the broad co-occurrence of stronger injury responses and lower rod relative abundance in the superior retina, rod abundance was not consistently different within focal response regions relative to the immediately flanking retina. Therefore, local variation in the response to degeneration occurred without corresponding differences in rod abundance at the timepoints examined. This does not exclude a relationship on a different timescale, as variegated patterns of photoreceptor light sensitivity and patches of photoreceptor loss have been reported in some human inherited retinal degenerations.^45,46^ Consistent with this possibility, several components of the rod injury response score used in this study have been shown to modify photoreceptor survival. Loss of leukemia inhibitory factor prevented the induction of all four components of the rod injury response score (*Edn2, Fgf2, Socs3,* and *Stat3*) and accelerated degeneration in the VPP retina.^32^ Similarly, increasing photoreceptor STAT3 or removing SOCS3-mediated feedback in rods preserved photoreceptors across genetically distinct models.^32,47,48^ Augmentation of FGF2 or *Edn2* can likewise increase photoreceptor survival, although the effects of endothelin signaling vary by disease model and intervention.^5,49,50^ Together, these perturbation studies indicate that pathways represented in the rod injury response can modify disease progression. Our findings add a spatial dimension by showing that expression of these pathway components is nonuniform across the degenerating retina. Longitudinal studies will be needed to determine how the focal response regions evolve over time and how they relate to subsequent photoreceptor loss or survival. We note that the mechanisms underlying these focal responses also need not be the same as those producing the broad asymmetry in rod transcriptional state and relative abundance.

Our cell type-resolved map further defines the cellular composition of the *LCA5^gt/gt^* retina in advanced disease. Prior work showed that the outer nuclear layer collapses to a single residual row of photoreceptor nuclei late in disease;^19^ we identify these surviving photoreceptors as predominantly cones. Although lebercilin localizes to the connecting cilium of both rods and cones,^17,18^ we show that cones persist substantially longer than rods in the *LCA5^gt/gt^* model. This finding provides a potential cellular basis for the relative sparing of cone function observed in individuals with *LCA5*-associated retinal degeneration,^19^ highlighting the translational relevance of this model. Notably, the persistence of cones provides important context for the improvements in cone-mediated vision reported in patients receiving LCA5 gene augmentation therapy in an ongoing phase 1b/2a clinical trial,^51^ suggesting that surviving cones may represent important targets for therapeutic rescue even at advanced stages of disease.

Several limitations should be considered. First, our 173-gene panel did not profile the full transcriptome and therefore may not have captured other degeneration-associated transcriptional changes with spatially patterned expression. The targeted panel design reflects a tradeoff between transcriptomic breadth and the single-cell resolution required to detect localized responses, such as those observed in rods. Higher-plex spatial profiling will be important for determining whether similar broad and focal patterns extend to other genes. Second, although the spatial patterns were concordant across two biological replicates, the small sample size and use of one model of degeneration limit assessment of the biological variability and generalizability of these patterns. Larger cohorts and additional models of inherited retinal degeneration will be required to determine whether similar spatial organization occurs in genetically distinct models of degeneration. Despite these limitations, our data reveal previously unresolved broad and focal patterning of degeneration-associated transcriptional responses and provide a framework for investigating whether anatomical position and local tissue context contribute to heterogeneity in photoreceptor susceptibility and therapeutic response, and, more broadly, neuronal vulnerability in neurodegenerative diseases.^52^

## Methods

### Mouse husbandry

*LCA5^gt/gt^* mice and wild-type C57BL/6J mice were housed at the University of Pennsylvania under a 12 h light/12 h dark cycle with food and water *ad libitum*. All procedures were performed in accordance with federal and institutional regulations under IACUC protocol #806931 and the ARVO Statement for the Use of Animals in Ophthalmic and Vision Research. Male mice were used for all conditions to limit sex-specific variation.

### Tissue collection and sectioning

Tissue collection was performed in mid-afternoon to reduce potential effects of the circadian rhythm. Mice were anesthetized by intraperitoneal injection of ketamine and xylazine and then perfused with 4% paraformaldehyde (PFA) in 1× PBS for 5 min. Eyes were enucleated immediately. Eyecups were then dissected in ice-cold 1× PBS, fixed in 4% PFA for 2 h at room temperature, washed for 1 h in 1× PBS at room temperature, cryoprotected overnight in 30% sucrose in 1× PBS at 4 °C, and embedded in OCT. Eyecup orientation was marked by visualizing the posterior ciliary arteries at the posterior surface of the eyecup, which mark the nasal-temporal axis.^53^ Eyecups were then flash-frozen in isopentane over dry ice. For sectioning, the OCT blocks were oriented using the nasal-temporal marks such that sections would span the superior-inferior axis. Once correctly oriented, 10 μm parasagittal sections were placed onto the Spatial Genomics coverslips. The sections were fixed using 4% PFA in 1× PBS for 10 min followed by three 5 min washes in 1× PBS and dried with 70% ethanol for 30 s before freezing at -80 °C.

### seqFISH

Coverslips were shipped to Spatial Genomics (Pasadena, CA 91107) on dry ice. seqFISH, DAPI and 18S rRNA staining, and transcript decoding were performed by Spatial Genomics (Pasadena, CA 91107) using the GenePS system and a custom 173 gene panel comprising 162 barcoded and 11 serially detected targets (Supplementary Table 1).

### Raw data handling

High-resolution images of DAPI and 18S rRNA staining and decoded transcript coordinates were imported from Spatial Genomics for analysis. Two manually drawn exclusion masks were applied to two regions containing overlapping or displaced retinal tissue caused by partial detachment from the slide. Because these regions could not be assigned confidently to the underlying retinal section, they were excluded from all analyses that followed cell segmentation, transcript assignment, and retinal layer annotation. Masks were drawn using the NimbusImage platform,^54^ which is available at https://github.com/arjunrajlaboratory/NimbusImage/ and hosted at https://www.nimbusimage.com/.

### Cell segmentation and transcript assignment

Cells were segmented using the pretrained cpsam model of Cellpose-SAM (v.4.1.1).^55^ Specifically, nuclei were first segmented using the DAPI channel. Then, cell bodies were segmented from the composite DAPI and 18S rRNA channels; the latter was normalized with contrast limited adaptive histogram equalization (CLAHE) to account for the large difference in 18S rRNA signal intensity across the retinal layers. When cell bodies contained multiple nuclei, they were split using watershed segmentation on the 18S rRNA channel and using the nuclei as seeds. When no cell body was detected for a given nuclear mask, the nuclear mask was used instead. Transcripts were assigned to a cell when their coordinates fell within that cell’s segmentation mask. To account for differences in cell size, expression of each gene was quantified as transcript count divided by the cell mask area in μm^2^. Segmentation coverage was quantified as the fraction of DAPI positive area contained within cell body masks (DAPI positivity was defined using an Otsu threshold computed per retinal section).

### Retinal layer annotation

To generate consistent retinal layer annotations across samples, we trained a five-class U-Net^56^ model on DAPI and 18S rRNA images that were downsampled eight-fold. Training labels were manually drawn on 18S rRNA images using the NimbusImage platform. 6 of the 7 annotated retinal sections were used for training and 1 section was held out for validation. The model was trained for 80 epochs using the AdamW optimizer and a combined cross-entropy and soft Dice loss. The version with the highest mean validation intersection-over-union (IoU) was selected. The model classified each pixel as background, outer nuclear layer (ONL), inner nuclear layer (INL), inner plexiform layer (IPL), and ganglion cell layer (GCL). The outer plexiform layer (OPL) was not annotated because it was too thin to delineate reliably.

### Cell type assignment

Cell types were assigned from canonical marker gene modules (Supplementary Table 2). For each retinal section, a marker gene module score was calculated for each candidate cell type using the sum of log_1_p-transformed raw marker gene transcript counts. Otsu thresholding was then used to define cells positive for a module. Cells were assigned to the cell type with the strongest z-scored module score across all cells in the retinal section. Cells that did not meet the threshold for any cell type were labeled unassigned and excluded from downstream analyses. Of 117,775 segmented cells not inside the exclusion masks, 16,434 were classified as unassigned and excluded, retaining 101,341 typed cells for downstream analyses.

### Orientation of retinal sections along the superior-inferior axis

The superior-inferior orientation of each section was inferred from the reciprocal cone opsin gradients of *Opn1mw* and *Opn1sw*. For the continuous superior-inferior gradient used to quantify the distribution of transcriptional responses in rods at 21 days (Fig. 3b) and to detrend the rod injury response score (Fig. 4), we calculated the difference between log_1_p-transformed *Opn1mw* and *Opn1sw* transcript counts in each cone and computed a Gaussian-weighted average (σ = 120 μm) over up to the 500 nearest cones at each position. For Supplementary Fig. 2c and Supplementary Fig. 4b, the smoothed cone opsin gradient was divided at the median for each retinal section, with higher values classified as superior and lower values as inferior.

### Dimensionality reduction

For visualization purposes only, typed cells from all conditions were embedded by UMAP^57^ after Harmony^58^ integration (v0.0.10). Expression profiles across all 173 genes were normalized to 10,000 transcripts per cell, log_1_p-transformed, and scaled using a z-score before performing principal component analysis. The first 30 principal components were integrated using Harmony with a maximum of 20 iterations and sample collection batch as the integration variable, and then embedded using UMAP with a k-nearest neighbors value of 15. Cell type assignment and downstream analyses were performed independently of the UMAP embedding.

### Cell type relative abundance and composition

For each retinal section, the relative abundance of each cell type was calculated as the number of cells assigned to that type divided by the total number of typed cells. Photoreceptor composition was calculated as the number of rods or cones divided by the total number of rods and cones. For Supplementary Fig. 2c, rod relative abundance was calculated separately within the superior and inferior hemispheres of each retinal section.

### Estimation of outer nuclear layer (ONL) and inner nuclear layer (INL) thicknesses

For each retinal section, DAPI positive pixels were identified using an Otsu threshold calculated from non-zero DAPI intensities. The DAPI pixels were then assigned to the ONL or INL using the corresponding U-Net layer annotations. An arc along the midline of each retinal section was computationally drawn separately for each contiguous piece of tissue within the section. For each layer, the DAPI positive area was divided by the total arc lengths of the individual tissue pieces.

### Total retina relative transcript abundance

For each retinal section, the relative abundance of each gene was calculated as the fraction of transcripts within the annotated retinal layers assigned to that gene. For each *LCA5^gt/gt^* replicate, the log₂-fold change was calculated relative to the mean relative abundance in the two age-matched wild-type replicates. A gene was classified as having changed when both *LCA5^gt/gt^*replicate log₂-fold changes were at least 0.5 in the same direction and both *LCA5^gt/gt^* values were either higher or lower than both wild-type values.

### Selection of genes with reliable rod expression

To reduce the influence of transcripts assigned to rod masks from neighboring cell spillover and/or extended cellular processes, genes were filtered using a relative rod expression score. To calculate the score for each gene, the mean transcript density was first calculated for each cell type after pooling cells from both *LCA5^gt/gt^* retinal sections at 21 days. The mean density in rods was then divided by the highest mean density observed across all cell types, including rods themselves. A score of 1 therefore indicated that rods had the highest mean expression of that gene compared to other cell types. Non-rod markers had scores no more than 0.103. Therefore, a minimum rod relative expression score of 0.15 was used as a threshold. Genes were additionally required to be detected at least once in 10% or more of rods in both replicates of at least one of the following conditions: wild-type 21 days, wild-type 64 days, *LCA5^gt/gt^* 21 days, or *LCA5^gt/gt^* 30 days. The *LCA5^gt/gt^* retinas at 64 days were not used for determining the detection criterion because very few rods remained. After applying these two filters, 56 genes were retained for the analyses in Fig. 3 and Fig. 4.

### Identification of rod transcriptional responses in *LCA5^gt/gt^* retinas at 21 days

For each retained rod gene, the mean rod transcript density was calculated separately for each retinal section. For each *LCA5^gt/gt^* replicate, the log₂-fold change was calculated relative to the mean rod transcript density in the two wild-type replicates. A gene was classified as having changed when both *LCA5^gt/gt^* replicate log₂-fold changes were at least 0.5 in the same direction and both mutant values were either higher or lower than both wild-type values.

### Superior-inferior distribution of rod transcriptional responses in *LCA5^gt/gt^*retinas at 21 days

For each gene classified as having changed in *LCA5^gt/gt^* rods at 21 days, a Spearman rank correlation was calculated separately for each *LCA5^gt/gt^* retinal section between each rod’s log_1_p-transformed transcript density and the smoothed cone opsin gradient at that rod’s position. Positive values indicated enrichment toward the superior retina, whereas negative values indicated enrichment toward the inferior retina.

### Construction of the rod injury response score

The rod injury response score was calculated using *Edn2*, *Socs3*, *Fgf2*, and *Stat3* expression. For each gene, log_1_p-transformed transcript density was standardized (z-scored) across rods pooled from all wild-type and *LCA5^gt/gt^*conditions. The four z-scores for each rod were then averaged with equal weight.

### Quantification of the number and size of focal rod injury response regions

Rods were projected onto the retinal arc for each contiguous piece of tissue and assigned to consecutive 25 μm columns along the arc. The mean injury response score of rods in columns containing at least one rod was calculated. To account for the broad superior-inferior gradient, a linear regression was fitted across scored columns between the mean injury response score and the mean smoothed cone opsin gradient within the same column for each retinal section. The predicted score was then subtracted from the observed score to obtain a residual injury response score. Columns with no rods were treated as gaps.

Within each continuous run of scored columns, the residual injury response profile was smoothed using a Gaussian smoothing width of σ = 37.5 μm. Smoothing was not performed across gaps or disconnected tissue pieces. Local maxima in the smoothed profile were then identified. The prominence of each local maximum was calculated as its height above the greater of its two local baseline minima. A focal region was defined as a local maximum with a prominence of at least 0.10. At the end of a scored column run, where only one side of the peak was observed, a peak was retained if it rose by at least 0.10 above both the minimum on its observed side and a residual injury response score of zero on the other (unobserved) side. Focal regions associated with these peaks were classified as ‘truncated’ but retained for analysis.

The boundaries of complete (not truncated) regions were defined where the smoothed profile declined halfway from the peak toward its local baseline. For truncated regions, the boundary was calculated on the observed side, and the end of the scored run was used as the boundary on the unobserved side. The distance between the boundaries was reported as the estimated width of the focal region. For truncated regions, this width necessarily represented only the observed portion of the region. When the half-prominence intervals of adjacent peaks overlapped, they were separated at the shared local minimum. A 25 μm column was assigned to a focal region when its center fell within these boundaries.

The sensitivity of the detection of focal regions was assessed by varying the peak prominence threshold from 0 to 0.50 while keeping the smoothing width at σ = 37.5 μm, and by varying the smoothing width from σ = 0 to 62.5 μm while keeping the prominence threshold at 0.10.

### Robustness of focal rod injury response pattern

To test whether the spatial organization of the rod injury response profile depended on any single score gene, each of the four genes was omitted in turn and the score was recalculated as the mean of the remaining three gene z-scores. For each *LCA5^gt/gt^* retinal section at 21 and 30 days, the three-gene score was detrended and smoothed using the same method described above, and its smoothed spatial profile was compared to the four-gene spatial profile using Pearson’s correlation.

### Rod abundance in focal injury response regions

For each focal region, the ratio of the mean number of rods per 25 μm column inside the region to the mean in immediately flanking reference columns was calculated. For complete regions, immediately flanking reference columns spanning up to one focal region width on each side were used. If either side overlapped another focal region, a column with no rods, or the end of a tissue piece, selection was stopped at that boundary and the reference on the opposite side was shortened to the same width. For truncated regions, up to one focal region width was used from the observed side.

### Rod transcript enrichment in focal rod injury response regions

To identify rod transcripts locally enriched within focal injury response regions, the mean transcript density among rods was calculated for each 25 μm column containing at least one rod. Within each retinal section, enrichment was calculated as the mean transcript density in focal region columns divided by the mean in the immediately flanking reference columns. The reference columns were selected using the method described above. A gene was classified as enriched when its enrichment was greater than 1 in all *LCA5^gt/gt^* retinal sections at 21 days and 30 days. Genes with a mean enrichment of at least 1.2 across these four sections were labeled in Fig. 4d. To avoid circularity in the calculation, the four injury response score genes (*Edn2, Socs3, Fgf2,* and *Stat3*) were excluded from the initial analysis. Each score gene was assessed separately by redefining the focal regions using the other three genes and then measuring enrichment of the omitted gene.

### Müller gliosis in focal rod injury response regions

To assess Müller gliosis marker enrichment in the focal regions, *Gfap* and *Serpina3n* transcripts in the ONL and INL were combined within each 25 μm column, then normalized to the combined ONL and INL layer area. Within each retinal section, enrichment was calculated as the mean combined transcript density in focal region columns divided by the mean in their immediately flanking reference columns. The reference columns were selected using the method described above. Columns containing no *Gfap* or *Serpina3n* transcripts were counted as zero. The overall Müller gliosis marker expression across wild-type and *LCA5^gt/gt^* retinas was calculated as the mean combined transcript density across the 25 μm columns. The number of Müller glia cell bodies (normalized to INL area) and the combined *Gfap* and *Serpina3n* transcript densities were correlated across columns within each retinal section using Spearman’s correlation.

### Microglia localization and abundance in focal rod injury response regions

The fraction of microglia localized to the outer nuclear layer was calculated separately for each retinal section. To assess enrichment in focal rod injury response regions, the mean number of outer nuclear layer microglia per 25 μm column within focal regions was divided by the mean number per column in the remaining analyzed retinal columns. Columns containing no outer nuclear layer microglia were counted as zero.

### Visualization of retinal sections and transcript density

For visualization of retinal sections only, DAPI and 18S rRNA images were scaled from the 2nd to 98th percentile and gamma-adjusted using γ = 0.55 and 0.7, respectively. In addition, color scales showing transcript density were scaled from zero to the largest 97th percentile value across the displayed retinal sections. Values outside of these display ranges were shown using the maximum corresponding color. For retinal sections colored by the rod injury response score, the same display method was used but instead scaled from the 2nd to the 98th percentile, as scores could be positive or negative. None of these transformations were used for quantitative analysis.

### Use of artificial intelligence tools

During the preparation of this work, the authors used Claude and ChatGPT to generate and improve code for analysis and suggest improvements to the clarity and flow of the text. The authors reviewed and edited the content and take full responsibility for the content of the manuscript.

## Data availability

Raw and preprocessed data can be found under the following link: https://doi.org/10.5281/zenodo.22767514

## Code availability

Code used to produce analyses and figures can be found under the following link: https://github.com/arjunrajlaboratory/retina_spatial_paper.git

## Acknowledgements

The authors thank Steven Chomistek for providing several mice, Brent A. Bell for advice on using retinal vasculature for orientation marking and providing the tungsten lamp, Dale L. Brokaw, Pavithran T. Ravindran, Catherine G. Triandafillou, Vinay Ayyappan, and Grant Kinsler for assistance with preliminary RNA FISH experiments and advice on the seqFISH workflow, and members of the Raj and Uyhazi labs for discussions and feedback on the manuscript.

## Author contributions

A.A.A. conceptualized the project and performed analyses under the supervision of A.R. and K.E.U. A.A.A. performed experiments with assistance from R.P. for tissue collection and Z.W. for tissue sectioning. S.K. contributed to the approach for quantifying focal region size and number. A.A.A. wrote the manuscript with supervision and editing from A.R. and K.E.U., and input from all authors. All authors read and approved the manuscript.

