## Supplementary Information for "Single-cell spatial transcriptomics of inherited retinal degeneration reveals broad and focal organization of disease responses"

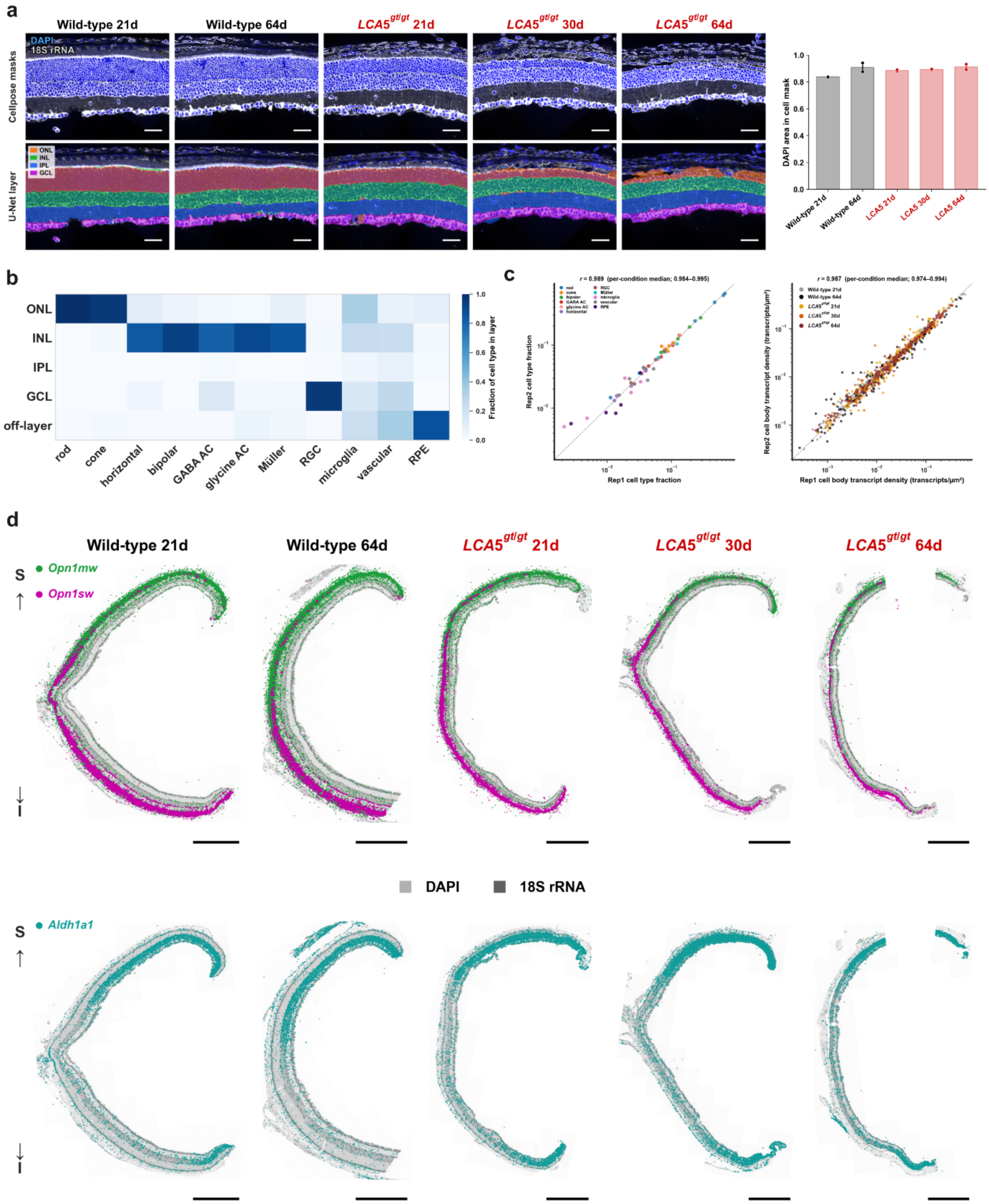


**Supplementary Fig. 1: Validation of cell segmentation, cell type assignment, and superior-inferior orientation. (a)** Left, top row, representative DAPI and 18S rRNA images with segmented cell masks for each condition. Left, bottom row, retinal layer annotations of the outer nuclear layer (ONL), inner nuclear layer (INL), inner plexiform layer (IPL), and ganglion cell layer (GCL). Scale bars, 50 μm. Right, fraction of DAPI signal captured within cell masks. Points represent individual replicates; bars show the mean; whiskers span the minimum and maximum replicate values. **(b)** Laminar distribution of each cell type, pooled across conditions and individual replicates. For each cell type, the color represents the fraction of cells located in the ONL, INL, IPL, GCL, or outside the annotated retinal layers ('off-layer'). **(c)** Pearson's correlation between individual replicates of each condition for $\log_{10}$-transformed values of the fraction of typed cells assigned to each cell type (left) and mean cellular transcript density for each gene (right). **(d)** Representative full retinal sections from each condition showing *Opn1mw* and *Opn1sw* transcripts (top) and *Aldh1a1* transcripts (bottom). Sections are oriented with the superior hemisphere at the top and the inferior hemisphere at the bottom. Scale bar, 500 μm.

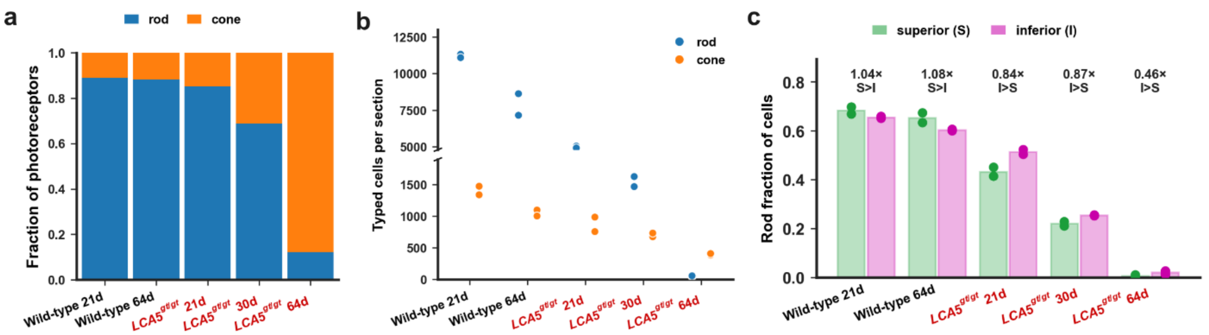


**Supplementary Fig. 2: Photoreceptor abundances across *LCA5*$^{gt/gt}$ degeneration. (a)** Fraction of photoreceptors assigned as rods or cones in each condition. Stacked bars show the mean of individual replicates. **(b)** Absolute number of rods and cones in each condition. Points represent individual replicates. **(c)** Fraction of total typed cells assigned as rods in the superior and inferior retinal hemispheres. Points represent individual replicates; bars represent replicate means; S>I or I>S labels indicate conditions in which both replicates agreed in direction.

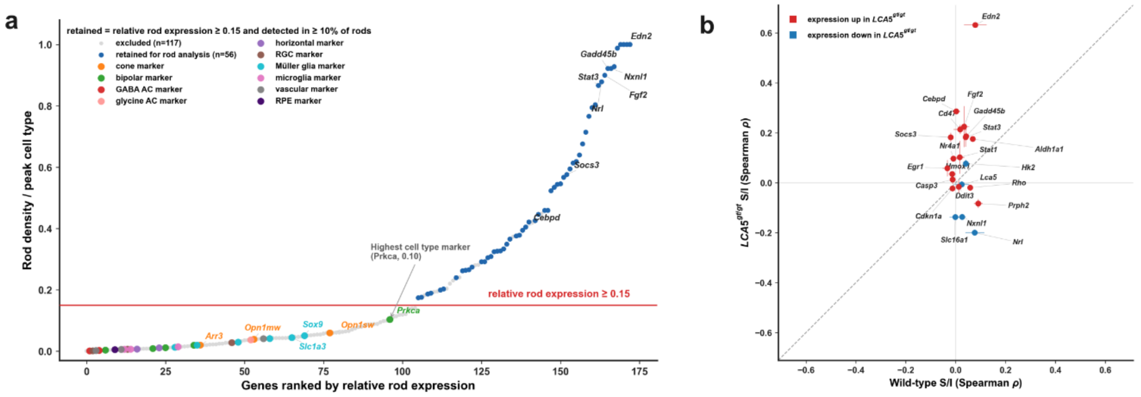


**Supplementary Fig. 3: Rod gene filtering and superior-inferior distribution of responses in wild-type vs *LCA5*$^{gt/gt}$ retinas at 21 days. (a)** Relative expression of all 173 panel genes in rods. Genes with a relative rod expression score ≥ 0.15 and detected in ≥10% of rods in both replicates of at least one condition were retained for analysis (blue; *n* = 56); genes below either threshold were excluded (gray; *n* = 117). **(b)** Superior-inferior distribution of the 22 genes in Fig. 3b in wild-type versus *LCA5*$^{gt/gt}$ retinas at 21 days, colored by the direction of change in expression (red = increased, blue = decreased). Each point represents the mean of individual replicates; horizontal and vertical bars span the two wild-type and *LCA5*$^{gt/gt}$ replicate values, respectively.

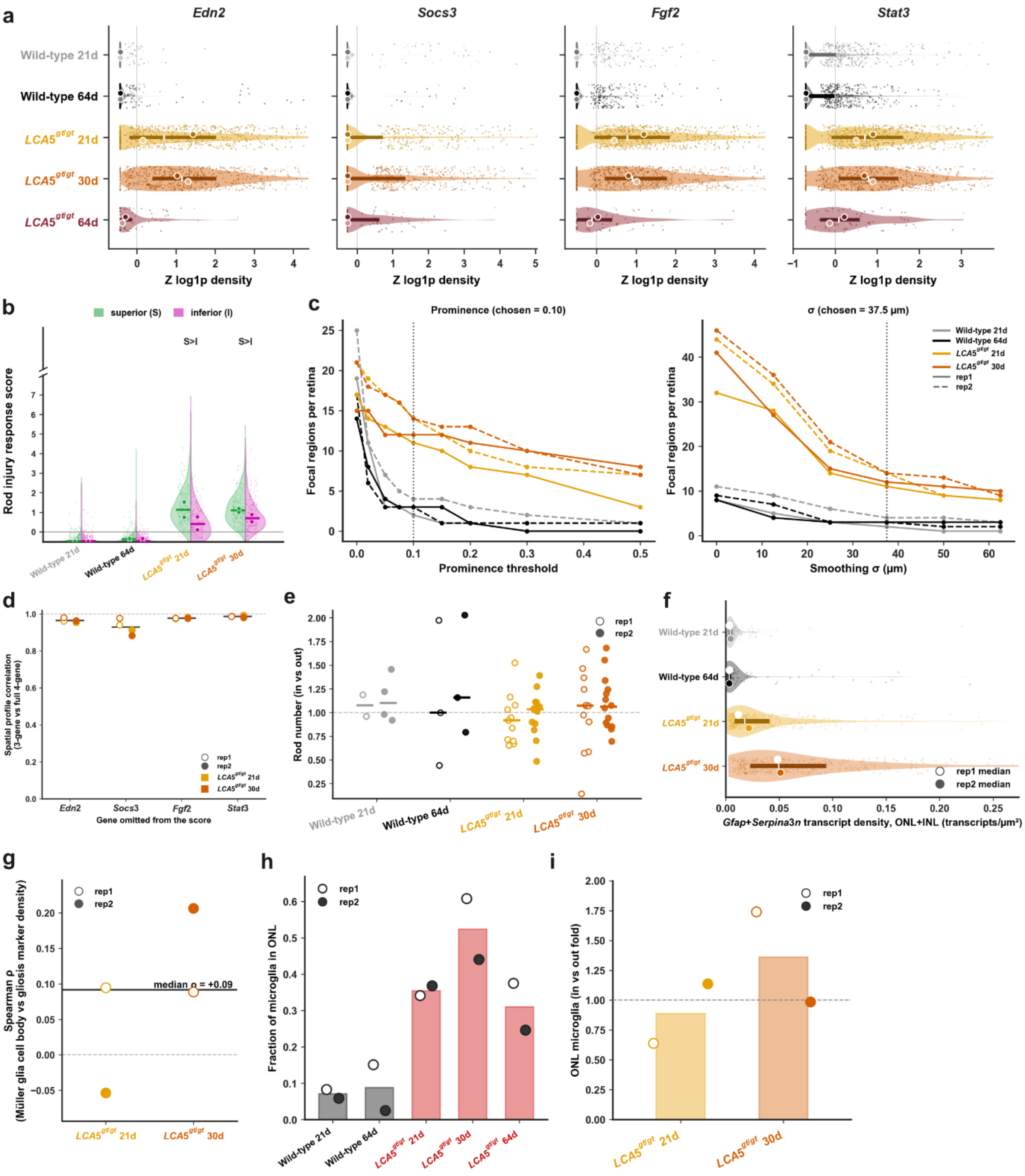


**Supplementary Fig. 4: Additional characterization of focal rod injury response regions.**

**(a)** Distributions of the injury response score components *Edn2, Socs3, Fgf2,* and *Stat3* in rods across conditions. Violins show all rods and points show a random subsample of up to 500 rods per replicate. Circles represent replicate medians; bars represent the pooled interquartile range;

white lines represent the pooled median. **(b)** Rod injury response scores in superior (S) and inferior (I) hemispheres of wild-type and $LCA5^{gt/gt}$ retinas. Horizontal lines indicate pooled medians and circles indicate replicate medians. S>I indicates conditions with a higher superior median in both replicates. **(c)** Sensitivity of focal region counts to peak prominence (left) and Gaussian smoothing width (right). Solid and dashed lines represent individual replicates. Vertical dotted lines indicate values chosen for analysis (prominence = 0.10; $\sigma$ = 37.5 μm). **(d)** Sensitivity of spatial organization of rod injury response to the exclusion of individual score genes. Circles show Pearson's correlations for individual retinal sections; horizontal lines show the mean across four sections. **(e)** Mean rod abundance in focal rod injury response regions relative to immediately flanking tissue in wild-type retinas at 21 and 64 days and $LCA5^{gt/gt}$ retinas at 21 and 30 days. Points represent individual focal regions; open and filled circles represent individual replicates. **(f)** Combined *Gfap* and *Serpina3n* transcript density in the outer and inner nuclear layers across wild-type and $LCA5^{gt/gt}$ retinas. Points represent 25 μm columns; circles represent replicate medians; bars represent the pooled interquartile range; white lines represent the pooled median. **(g)** Spearman correlation between Müller cell body density and combined *Gfap* and *Serpina3n* transcript density in $LCA5^{gt/gt}$ retinas at 21 and 30 days. **(h)** Fraction of microglia localized to the outer nuclear layer in all conditions. Bars show the mean of two replicates; open and filled circles represent individual replicates. **(i)** Microglia counts per 25 μm column in the outer nuclear layer within focal regions relative to the remaining outer nuclear layer in $LCA5^{gt/gt}$ retinas at 21 and 30 days. Bars show the mean of two replicates; open and filled circles represent individual replicates.

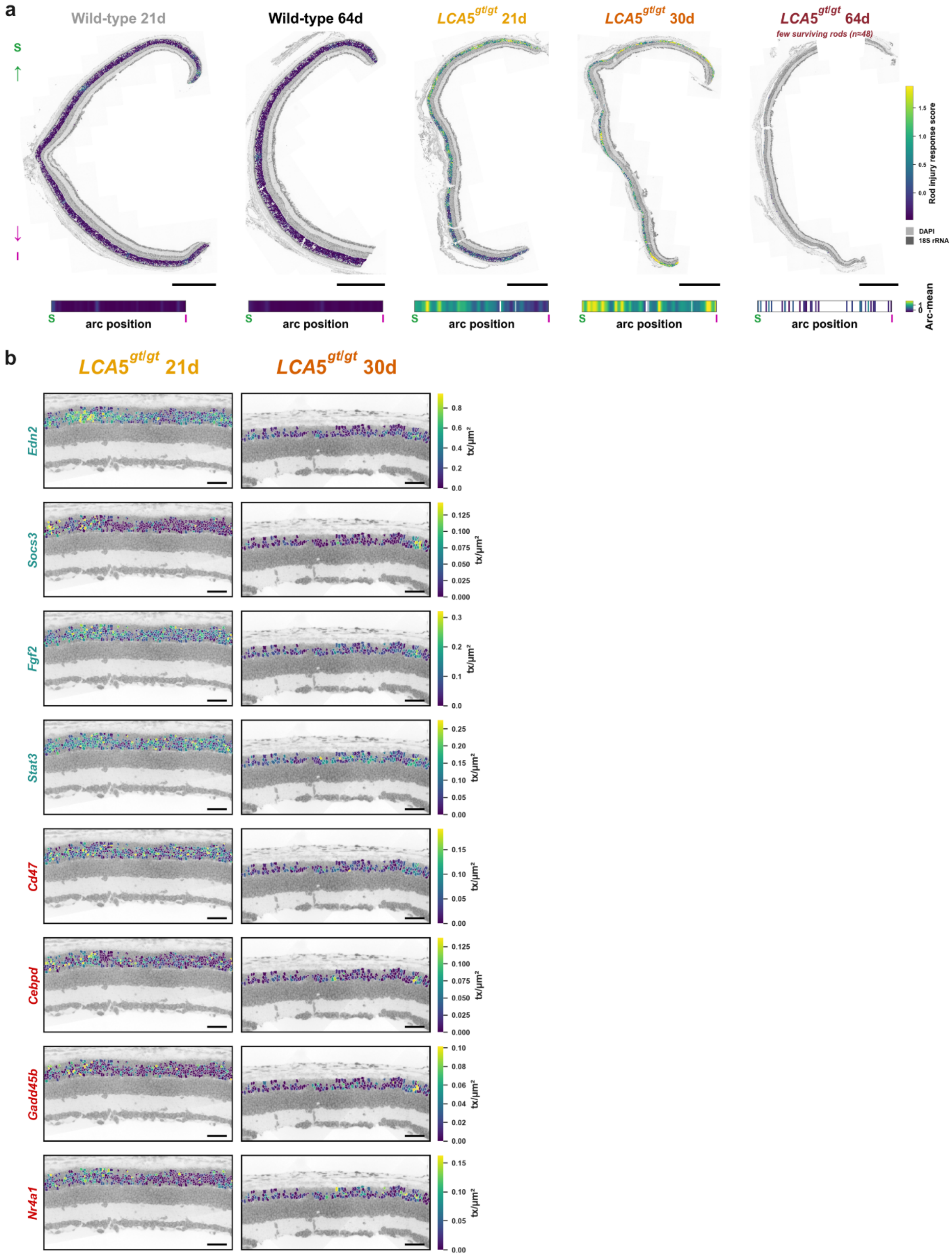
a
Wild-type 21d
Wild-type 64d
LCA5gt/gt 21d
LCA5gt/gt 30d
LCA5gt/gt 64d
few surviving rods (n=48)
S
I
Rod injury response score
DAPI
18S rRNA
arc position
Arc-mean
b
LCA5gt/gt 21d
LCA5gt/gt 30d
Edn2
Socs3
Fgf2
Stat3
Cd47
Cebpd
Gadd45b
Nr4a1
tx/µm²

**Supplementary Fig. 5: Focal rod transcriptional responses across conditions and individual genes. (a)** Rod injury response score mapped onto representative sections from all conditions. Horizontal strips show mean rod injury response score in 25 µm columns along the retinal arc. **(b)** Magnified regions of *LCA5*$^{gt/gt}$ retinal sections at 21 and 30 days showing the rod transcript density of *Edn2*, *Socs3*, *Fgf2*, *Stat3*, *Cd47*, *Cebpd*, *Gadd45b*, and *Nr4a1*. Each gene is shown on its own scale. Scale bars, 50 µm.

**Supplementary Table 1:** List of 173 genes in seqFISH panel sorted by detection method

| **Detection method** | **Genes** |
|---|---|
| Barcode (162) | A2m, Acan, Acsl4, Aldh1a1, Aqp4, Ascl1, Atf4, Axin2, Bax, Bcl2l1, Bsn, C1ql2, C1ql3, C3, C3ar1, Calb1, Capn2, Casp1, Casp3, Ccl2, Ccn2, Ccr2, Cd44, Cd47, Cd68, Cd74, Cdh2, Cdkn1a, Cebpd, Cldn5, Cntf, Crb1, Crb2, Crx, Csf1r, Ctbp2, Cx3cl1, Cx3cr1, Cxcl10, Cyp26a1, Dab1, Dag1, Ddit3, Edn2, Ednrb, Egflam, Egr1, Elfn1, Fgf2, Fn1, Gad1, Gadd45b, Gap43, Gas6, Gfap, Gja1, Gjd2, Gpnmb, Grm6, Gsdmd, Hes1, Hk2, Hmox1, Hspa5, Ift88, Igf1, Igf1r, Il10, Il10ra, Il1b, Il1r1, Il34, Irf7, Irf8, Isl1, Itgam, Itgax, Kcnj10, Lca5, Lcn2, Lgals3, Lif, Lifr, Lpl, Map1lc3b, Mdk, Megf10, Mertk, Mki67, Mlkl, Mmp9, Ncan, Nefh, Nes, Nfkbia, Nlgn2, Nr2e3, Nr4a1, Nrl, Nrp1, Nxnl1, Onecut2, Opn4, Osmr, Otx2, P2ry12, Pax6, Pecam1, Pou4f2, Prom1, Prox1, Ptn, Ptprs, Ptprz1, Rara, Rbpms, Rgs5, Rims1, Ripk3, Rpe65, Scgn, Sdha, Sema3a, Serpina3n, Serpinf1, Serping1, Sirpa, Slc16a1, Slc16a3, Slc17a7, Slc1a3, Slc2a1, Slc32a1, Slc6a9, Slc7a11, Socs3, Sod2, Sox2, Sox9, Spp1, Sqstm1, Stat1, Stat3, Tgfb1, Tgfbr2, Thy1, Timp1, Tjp1, Tmem119, Tnc, Tnf, Tnfrsf1a, Trem2, Txnip, Tyrobp, Vcan, Vegfa, Vim, Vsx1, Vsx2, Xbp1, Xiap |
| Serial (11) | Apoe, Arr3, B2m, C1qa, Glul, Opn1mw, Opn1sw, Pde6b, Prkca, Prph2, Rho |

**Supplementary Table 2:** Marker genes used to assign 11 major cell types

| **Cell type** | **Marker genes** |
|---|---|
| Rod | Nrl, Nr2e3, Pde6b |
| Cone | Arr3, Opn1mw, Opn1sw |
| Bipolar cell | Vsx2 alone, or Vsx2 with one of Prkca, Grm6, Vsx1, or Scgn |
| GABAergic amacrine cell | Gad1, Slc32a1 |
| Glycinergic amacrine cell | Slc6a9 |
| Horizontal cell | Onecut2, Calb1 |
| Retinal ganglion cell | Rbpms, Pou4f2, Nefh |
| Müller glia | Sox9, Slc1a3, Glul, Aqp4 |
| Microglia | Csf1r, Cx3cr1, C1qa, Ccr2 negative |
| Vascular cell | Cldn5, Pecam1, Rgs5 |
| Retinal pigment epithelium | Rpe65 |
